# survInTime - Exploring surveillance methods and data analysis on Brazilian respiratory syndrome dataset and community mobility changes

**DOI:** 10.1101/2023.09.26.559599

**Authors:** Yasmmin Côrtes Martins, Ronaldo Francisco da Silva

## Abstract

**Background:** The covid-19 pandemic brought negative impacts in almost every country in the world. These impacts were observed mainly in the public health sphere, with a rapid raise and spread of the disease and failed attempts to restrain it while there was no treatment. However, in developing countries, the impacts were severe in other aspects such as the intensification of social inequality, poverty and food insecurity. Specifically in Brazil, the miscommunication among the government layers conducted the control measures to a complete chaos in a country of continental dimensions. Brazil made an effort to register granular informative data about the case reports and their outcomes, while this data is available and can be consumed freely, there are issues concerning the integrity and inconsistencies between the real number of cases and the number of notifications in this dataset.

**Results:** We projected and implemented four types of analysis to explore the Brazilian public dataset of Severe Acute Respiratory Syndrome (srag dataset) notifications and the google dataset of community mobility change (mobility dataset). These analysis provides some diagnosis of data integration issues and strategies to integrate data and experimentation of surveillance analysis. The first type of analysis aims at describing and exploring the data contained in both datasets, starting by assessing the data quality concerning missing data, then summarizing the patterns found in this datasets. The Second type concerns an statistical experiment to estimate the cases from mobility patterns organized in periods of time. We also developed, as the third analysis type, an algorithm to help the understanding of the disease waves by detecting them and compare the time periods across the cities. Lastly, we build time series datasets considering deaths, overall cases and residential mobility change in regular time periods and used as features to group cities with similar behavior.

**Conclusion:** The exploratory data analysis showed the under representation of covid-19 cases in many small cities in Brazil that were absent in the srag dataset or with a number of cases very low than real projections. We also assessed the availability of data for the Brazilian cities in the mobility dataset in each state, finding out that not all the states were represented and the best coverage occurred in Rio de Janeiro state. We compared the capacity of place categories mobility change combination on estimating the number of cases measuring the errors and identifying the best components in mobility that could affect the cases. In order to target specific strategies for groups of cities, we compared strategies to cluster cities that obtained similar outcomes behavior along the time, highlighting the divergence on handling the disease.

**Availability:** https://github.com/YasCoMa/dashboard-srag-mobility

## 1 Introduction

In Brazil, the first case of covid-19 was reported in February 26, 2020 in São Paulo city [1], one of the largest capitals with a high population density containing 12,200,180 inhabitants (according to the 2022 census results^1^). The airports were not immediately closed and the control measures at the time were not implemented efficiently even seeing the projections and the death track coming from the European countries and Asia. The latency to respond [2] to the disease arrival by restricting the people mobility or encouraging hygiene measures such as mask usage and hand sanitizer cost the lives of more than 700,000 citizens. Brazil has a powerful free health system that every citizen and foreign people can be treated according to the health units and hospitals vacancies distributed along the cities in the 27 states. But this system has some bottlenecks and before this respiratory disease outbreak, it was already suffering from overcrowding in large cities for recurrent diseases and surgeries.

In order to organize the cases notification in the cities across the country, Brazil maintained a database (srag) of severe acute respiratory syndrome reports^2^, where the rows represent each patient information without identification details and the columns represent types of symptoms, risk factors, pregnancy status, gender, race, notification/residence state and city, etc. This database contains more than 3 million records but not all cities are represented in it neither the most part of the asymptomatic cases. The epidemiological works [3–10] published about the disease spread and cases simulation and projection clearly demonstrated the discrepancies between the data reported and the real number of cases and deaths across the Brazilian cities. However, few works provided integration and methodological approaches to diagnose and extract useful insights from this data. During the pandemic Google provided reports^3^ concerning the mobility percentile change across cities of many countries, showing mainly the reduction of tracked devices connected in outside places where the lockdown was successfully implemented. Recent and broad literature published about covid-19 in Brazil have not explored the combinations of data derived by both datasets to reveal epidemiological correlations.

In this paper, we aim at presenting four analytical methods to get useful insights from these data and learn lessons that can be used in a future pandemic of some disease with a similar spread profile. These methodological approaches are: (i) enable n exploratory data analysis from the data processing and cleaning to visualization resources with filters to extract insights from the available dimensions of data in each dataset, as well as diagnosing dataset issues concerning their quality; (ii) combine mobility change categories to estimate the normalized number of cases to answer the role and impacts of mobility change in the disease spread; (iii) detection and comparison of waves in multiple time series to understand the dynamics in diverse levels of population density across the Brazilian cities; finally, (iv) present two strategies to group cities based on time-dependent dimensions (cases, deaths, residential change) behavior, one of them based on graph communities [11] and the other using traditional machine learning algorithm (KMeans) [12].

## 2 Methods

### 2.1 Datasets

In this paper, we used two datasets, the first one was the Severe Acute Respiratory Syndrome dataset provided by the brazilian government that tracked information of each individual record of any respiratory disease case, including those that were confirmed to be caused by COVID-19^4^. The second dataset^5^ is the community mobility reports provided by Google, according to the usage of devices in certain place categories.

#### 2.1.1 Brazilian covid-19 notification report dataset

This dataset (srag) is released by year, we took into consideration the versions of 2020 and 2021. The 2021 edition added extra columns to handle the information about the vaccination like the vaccine manufacturer, doses, batches, etc. The dataset in 2020 had 153 columns and 13 vaccine related columns were added. In all the analysis showed in this paper, we do not considered any vaccine related information, we focused on the distribution of cases and their outcomes (cure or death) that were confirmed by PCR exam test [13], along some important patient characteristics such as birth date, notification city and state, residence state and city, gender, race, pregnancy stage (if applicable), risk factors, date of first symptoms, date of outcome decision and date of notification. These are the columns given by the raw files but we transformed the contents of these columns to normalize, correct formats and estimate derived column metrics.

The srag dataset was filtered according to the rows that contained results of PCR test positive for the SARS-CoV-2 virus. In order to guarantee the correct integration with the mobility change dataset, another filtering step removed the rows with residence and notification cities empty or that did not match the correct names registered in the last brazilian census. The Brazilian Institute of Geography and Statistics recently released the update population number of 5570 cities according to the 2022 census^6^. We integrated the brazilian states information^7^ together with the respective cities and their population to facilitate analysis and curate the residence and notification city fields.

The notification, birth, first symptoms and outcome decision dates were filled in the brazilian format, and in some cases were not complete with day, month and year. We tested the field and converted to the standard year-month-day format (yyyy-mm-dd), so that we could use to calculate other columns. Using these date columns we calculated the delay from the emergence of the symptoms till the notification, the delay from the notification to the outcome decision, and the age of the patient at the time of the notification. We classified the record using the age column according to the following age groups: 0-5, 6-12, 13-20, 21-40, 41-70, 71-120. We created a column to indicate the displacement of people for medical care, which is a binary field according to the difference or not between the notification and residence cities. From the notification date we derived the corresponding week and month by year to facilitate the stratification of time series analysis.

The gender, pregnancy status, outcome and race columns used numerical codes in the raw dataset, we retrieved the data dictionary to map back to the original values translated to english. The outcomes could be Cure or Death, and the sum of records in these two classes refers to the total number of cases. Concerning the risk factors, there are eleven columns filled as binary values for the following risk factors: heart disease, hematology, down syndrome, liver disease, asthma, diabetes, neurological disease, lung disease, immunosuppressive, kidney disease and obesity. But since there is another free text column for other types of risk factors, in which the other values were separated by semicolon. We united the standard eleven risk factors, when present, with those pointed in this free field, maintaining the separation by semicolon.

#### 2.1.2 Community mobility change dataset

This dataset (mobility) provides insights retrieved by the people devices with enabled place tracking feature, that allows the capture of the movement in certain places registered in google maps. In 2020, Google freely released an aggregated analysis comparing the mobility flow before pandemics (named as baseline) to the new numbers obtained for the same places organized by their categories, which are retail and recreation, groceries and pharmacy, transit stations, parks, workplaces, and residential. Although this dataset has some data quality issues such as the absence of data from people that do not have a smartphone or have it but turned off this tracking functionality.

In Brazil, many small cities are not contained in the dataset, in others some of these categories are missing. From the 5570 cities in Brazil 2254 have data in this dataset and 2214 have some column missing information. For this dataset, the values for the six categories are given as percentages, that may be negative (reduction) or positive (increase) in relation to the baseline. We maintained the date as it was already in the standard format, but we derived the week and month representation to perform aggregated time series analysis together with the data in the srag dataset. We also treated the state information, remove the prefix *State of* and converting the entire name (ex.: Rio de Janeiro) to the respective federative unit abbreviation (RJ), since the srag dataset uses this code for the state.

From these two final processed datasets, we created a third dataset combining the srag cases and outcomes according to the week and month periods of time, using these periods and the city name to integrate their data. We took the mean of the place categories percentage in the mobility dataset and used the count aggregation function to group the rows by period in the srag dataset. From these count, we also normalized by the city population and taking the number for each 1000 individuals, and the last derived feature is this count per 1000 in log10 scale.

### 2.2 Datasets exploration dashboard

We developed a web tool dashboard^8^ (Figure 1) to explore and filter the processed datasets using intuitive visualization plots. The dashboard is composed by three main pages, two of them provides insights and filtering options to query data quality metrics and specific insights about each dataset. The third uses the time series dataset combining data from srag and mobility to create a network composed by city nodes, whose edges are given by the euclidean distance of the numerical features representing them.

**Fig. 1.**
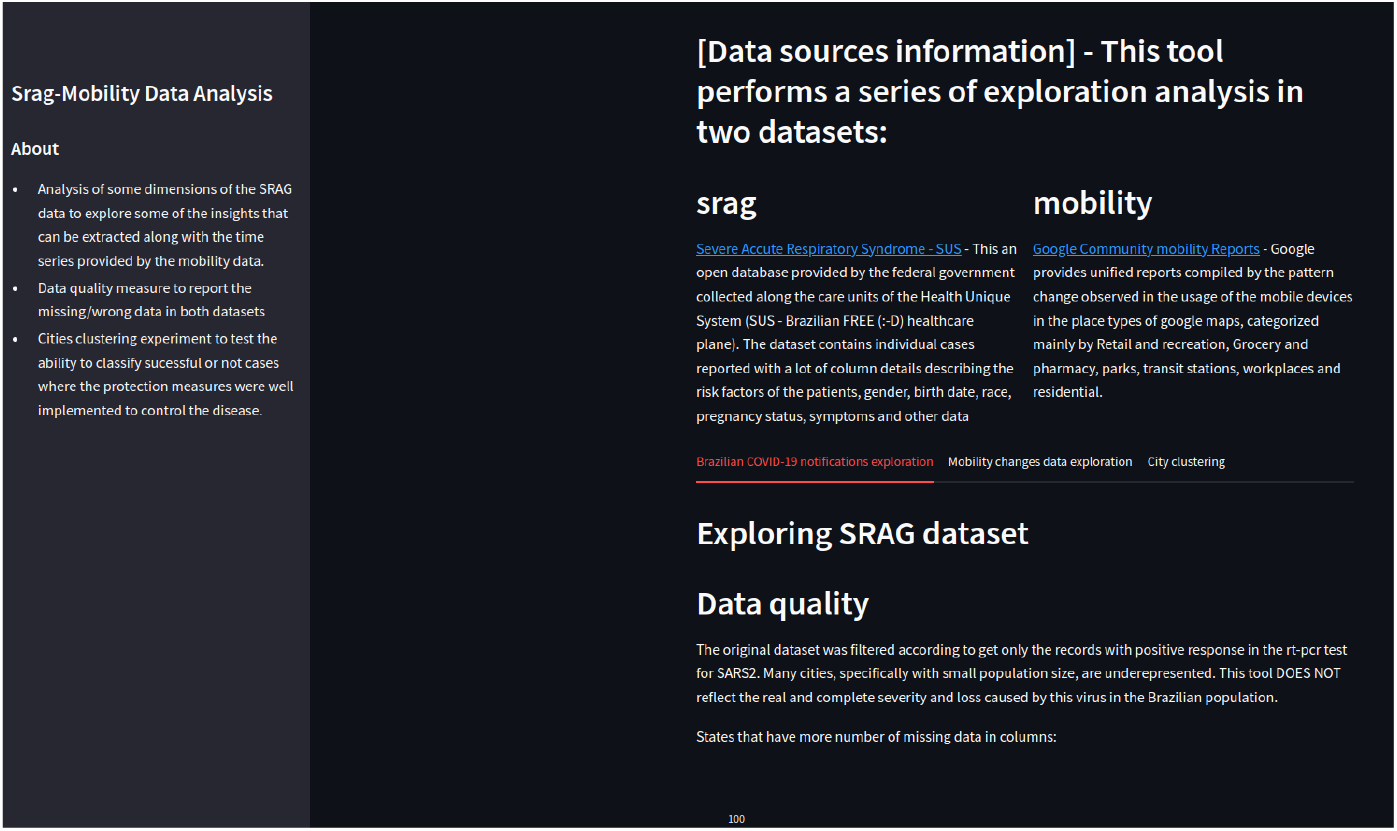
Screenshot of the survInTime web tool dashboard.

#### 2.2.1 Analysis of srag dataset

In the srag dataset, the first analysis section reports the missing data metrics such as the mean percentage of rows per state considering all columns that have empty values for some record. To demonstrate the specific columns that are contribution to these portions, it is possible to explore the percentage of rows with empty values for each specific column given a brazilian state of interest. The last section of the data quality analysis show the number of removed rows due to a mismatch between the city names in the srag dataset and the correct names listed in the most recent version of the brazilian census in 2022.

The next analysis section shows the aggregated count of notifications for each outcome (death and cure), all cases and also the number of cases in which the patient had to travel from its residence city to get health assistance in another larger city, that may be because of more vacancies or possibility to perform a more reliable diagnosis exam. We organized the results of this section according to the capital cities of each state. The counts were normalized by the population size per 100 to turn the results comparable.

The third analysis section allows the user see the geographical distribution of the scope desired (all cases, deaths or cure) in a specific state to get a clear cities border separation. The chosen scope is mapped according to a color scale that goes from heavy blue (low values) to light yellow (high values). All the other visualizations available also allow the filtering by state.

The fourth analysis section is oriented in distributing the results along time periods (weekly or monthly). We designed an interactive plot to compare the spread of new cities reporting the cases across the periods. This analysis demonstrates the speed and the delay of the virus arrival across the distinct states. Mainly from the economic and touristic hubs and the rest of the states. We allow the user to compare the number of new cities in the time line for up to four states. Based on the chosen type of time period (by month or week), the interactive dashboard allows the exploration of the scope dimension count across a diversified range of the feature columns available in the processed srag dataset: multiple pregnancy status that are first, second and third quarter; the types of risk factor found in the fixed and free text fields; gender; race; age groups; and just the existence or not of any risk factor. In the columns that have many feature values (ex.: types of risk factor), we selected the ten most popular ones in the dataset.

The last analysis section is focused on the top 15 ranked cities of the chosen state and dimension according to the records distributed according to the following feature columns in each plot: stratified types of risk factor; existence or not of risk factor; the delay in number of days taken between the first symptoms and the notification date; the delay also in days between the notification the decision outcome; and finally the comparison among the number of records that involved travel or not to seek healthcare in another city.

#### 2.2.2 Analysis of mobility dataset

The first analysis section for this dataset also evaluates aspects of its data quality. The first visualization plot is calculated based on the top 10 states with the highest number of cities covered by the community mobility change report. In a similar manner, we also measured the the mean percentage of missing data in each state considering the place categories; it is also possible to visualize the percentage of each of the six place categories percentage by choosing a specific state.

The second analysis section used the time series derived dataset to show the correlation among the place categories percentage and the three types of dimension scope of outcomes (overall cases, death or cure) normalized by the cities population per 1000. The user may filter according to the type of time period (week or month) and a specific state. We chose to force the filter per state capital city because the behavior varied a lot in each city due to a chaotic restriction policy implementation. This visualization resources provides a heatmap showing the strong/weak negative/positive correlation among the outcomes and place categories values.

To demonstrate the diversified distribution of the place categories percentages along the cities, the third analysis section compares the residential category with a second place category chosen by the user using a box plot formed by the values of these categories for all cities of each brazilian state. The existence of outliers in all states and the variations of median values in the categories are responsible to illustrate aforementioned chaotic behavior.

The last analysis section uses the time series dataset, and shows only the residential percentage change (primary y axis) and the overall cases normalized per 1000 (secondary y axis) aggregated according to the mean of the city values of a chosen state. The user can also choose the time period type (week or month) and the order priority (low to high, or the inverse direction). The x axis shows these values according to the top 15 cities in the state ordered following the mentioned filters criteria.

#### 2.2.3 City clustering

The last part of the dashboard aims at finding a pattern among the behavior observed across the periods of time to group all the cities that exist in both mobility and srag datasets. Based on the time series derived dataset integrating the outcome counts and the place categories percentage that co-occur in the same weeks and months, we generated a new dataset in which the columns are formed by all weeks of 2020 and 2021 and the rows are the cities. The values in each cell of the matrix are the aggregated mean of the chosen dimension scope (residential percentage change, deaths or overall cases). In case a city did not have data for a specific time period this column received zero.

Using this dataset, we formed a network [11] where the nodes are the cities and the edge among these nodes is calculated according to the euclidean distance of their numerical features. The user may choose a cutoff value to prune the edges indicating a high distance among the cities (default is 10). The capital cities are showed in the graph as blue nodes and the other ones are represented as green. We also provide to the user a list of the top 20 hub [14] cities that concentrate more connections with other cities and we also analyzed the community detection in this network and showed for each hub the number of communities that they participated.

### 2.3 Cases estimation from mobility change features

Although the mobility dataset has some completeness issues by not covering the population that do not have a device to track or that have but does not enable the GPS service or wireless connection, we considered the possibility of estimating the outcome scenarios and cases from the percentages of the six place categories. For all cities we performed a linear regression experiment to fit a model from the mobility features to explain the cases, cure and death outcomes.

The most important place category to detect lockdown and movement restriction is the residential, that should be negatively correlated with all the other five categories that indicate public/private places outside home. So, we tested in the model formula the combination of this variable alone and added with one of the other categories, in two estimation methods, which are ordinary least squares (ols) [15] and generalized linear models (glm) [16]. The performance was measured according the residual results provided by each method and the calculation of the root mean squared error (mse) [17].

### 2.4 Waves detection and comparison

Using the outcomes trajectory along the time periods, we designed and implemented an algorithm to detect the waves of the disease across the overall cases time series of all cities in the srag dataset in 2020 and 2021. These waves are characterized in disease epidemiology by time intervals where, in the case of a highly contagious viral infection, a new variant accelerates the transmission rate and new cases are reported and has the potential of causing many deaths. As the epidemiological models predict, this wave reaches a peak and the new cases start to decrease, since the number of susceptible people decreases [18].

The algorithm takes as input the times and the cases count in each time aggregated by week or month, and also the window interval to consider the found patterns a wave, so the minimum tiem period of a wave is two times the window value. The time series is then ordered from the oldest to the latest and at the first step it passes through the values marking the indexes and the trend (up or down) in each time step, by comparing the the next value with he current.

In the second part of the algorithm, it monitors the changes in the trends, adding the indexes in a temporary list of period times while the tend keeps increasing (up). In some cases, there is a little decrease to follow the rise again so we check the next neighbors and if those increase the indexes keep being stored. The algorithm stores the indexes of the wave candidates and it also tests the peak of the wave and keeps storing the periods of a same wave till finding another rising pattern. Once the it reaches the end of the observable period or the rising of a new potential wave, it tests if it has the minimal window as fixed in the cutoff and adds to the wave list.

We ran this algorithm and compared the overlapping of these waves along the cities to check the contagious pattern that occurred in the individual cities. We also Analyzed duration of the waves and number of drifts achieved in the cities.

### 2.5 Cities clustering by behavior pattern

In this analysis, we projected an experiment that takes two methodologies in consideration to group the brazilian cities with similar behavior pattern during the pandemic. We used the same derived and transposed dataset as presented in section 2.2.3, with the columns being a scope dimension (cases, deaths or residential percentage) per week time period, and the rows representing the cities. The first clustering method was the community identification by connected component detection, using the networkx [19, 20] in a graph formed by the cities as nodes and the euclidean distances among these cities as edges. In the web tool, the user may set a cutoff to prune the edges, but for this analysis we only consider the distance of 10.

The second method to group the cities is based on the traditional machine learning approach KMeans to tackle unsupervised learning [21]. In order to define the best number of groups, we applied the elbow method [12], then the clustering algorithm was executed varying the number groups (k) from 2 to 30 and measured the inertia to compare with the previous experiment using k-1 groups. We stopped the loop when the inertia reduction ratio ((previous inertia - current inertia)/previous inertia) reached the cutoff value of 0.01. We generated a report with the best k found for each scenario, the cities in each group and the total number of elements in it.

The main goal of this clustering analysis is comparing the agreement of city labeling in each clustering approach, and also checking the intersection of neighbors of the cities using the same scope dimension but with distinct clustering methods.

## 3 Results

The survInTime dashboard has the goal of describing the data contained in the srag and mobility datasets, and it does not aim at providing epidemiological analysis and trends. There is a vast literature about covid-19 using time series of cases and deaths to perform forecasting and reproduce complex infection modeling. The approaches presented in this paper are oriented to address the following research questions: (i) Explore new variables and distribution of their values time and city-oriented in srag dataset; (ii) demonstrate the chaotic distribution of mobility change in brazilian cities; (iii) assess the performance of mobility change in place categories on predicting the overall cases; (iv) detection of time intervals corresponding to waves from numerical series of cases; (v) find behavior patterns through cases, deaths and residential percentage change aggregated by week intervals.

### 3.1 Exploration of the srag dataset

There was a great effort in the entire country to concentrate and document in details the covid-19 cases together with many important variables such as the patient profile (race, gender, age, etc) and health prior conditions like post-partum situation, pregnancy stage, presence of risk factors, the symptoms reported, etc. In general, the raw dataset has 3,286,359 records considering 2020, 2021 and 2022. Only a fraction of these records (1,179,821) passed the filters of correct city names and confirmed cases by the gold standard PCR test. Besides these records have complete and curated information, they refer to only 3.6% of the 33,076,079^9^ cases of covid-19 in Brazil up to July 15 in 2022.

According to the missing data evaluation, most states obtained less than 15% of rows with some column ignored or empty. The states of Acre and Distrito Federal obtained the highest percentages (40.02 and 23.91, respectively). In these states, the most missing column is the *date outcome, pregnancy* (considering the female records that answered pregnancy status but ignored the stage) and *race*. A total of 1102 records were removed because due to a misspelling of the residence e/or notification city, that would not match to the cities reported in the mobility dataset in the data integration. More specifically, two cities corresponded to 85% of these records, which are *Sant’Ana do Livramento* (348 records) and *Mogi Mirim* (590). The names that were in the records for these cities had few typos: *Santana do Livramento* (Rio Grande do Sul) e *Moji Mirim* (São Paulo).

The summary of all cases (Figure 2) and the distinct outcomes (cure and death) in the state capitals show that the trend in the number of people that dislocated from their city of residence to seek health assistance in other city is similar to the overall cases. Many small cities initially did not have equipment to collect material to run the PCR test or did not have vacancies in the health basic units. The unified health system in Brasil allocates these basic units in all cities, but they have limited resources for emergence, when the patient needs some complex intervention, they are transferred to the closest reference hospital. So it is expected these counts for dislocation in the capitals. The overall cases normalized by the incidence of cases per 100 individuals in these cities indicate that Curitiba, Belo Horizonte, Recife, Porto Alegre e São Paulo obtained the highest number of case notifications. Interestingly, the data records show that all notifications in Brasília are from people from other city, these mistakes happened because the values correspond to administrative regions around Brasília, they were filled in the form as other regions but according to the census they are still considered as in Brasília city.

**Fig. 2.**
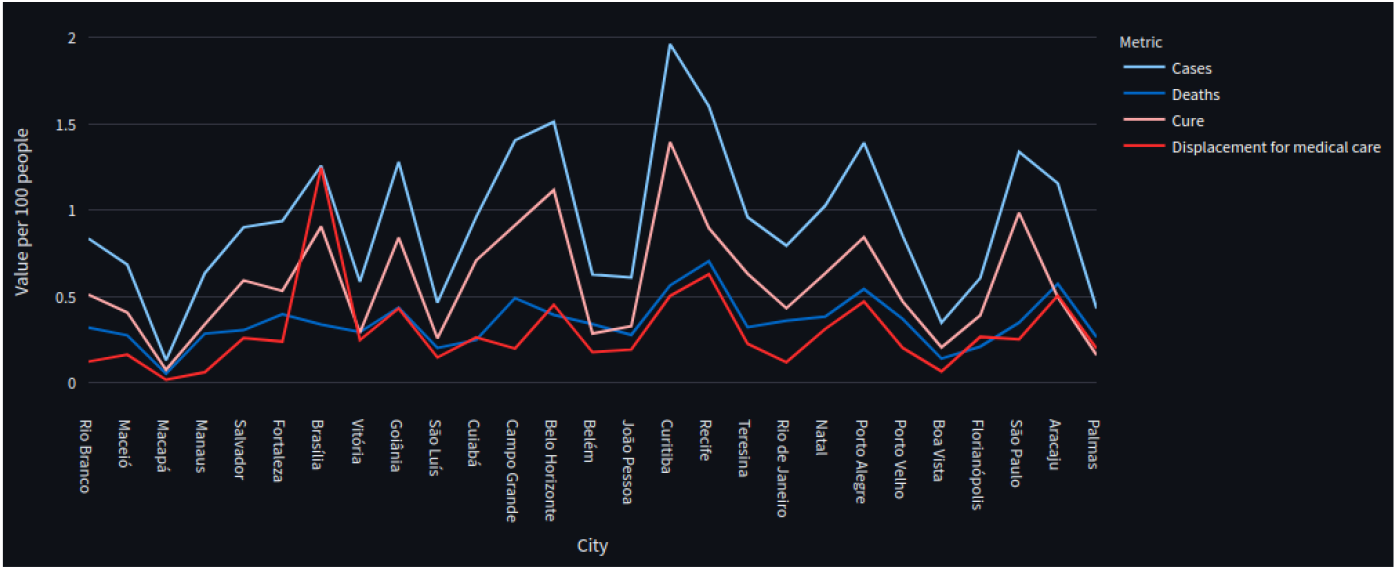
Comparison of overall cases, deaths, cured and displacement counts along the capital cities.

The comparison of the number of new cities that reported the disease along the weeks of 2020 and 2021 show that the disease arrived in the cities around the São Paulo capital faster than the other states we compared (Rio de Janeiro, Minas Gerais and Ceará). In Minas Gerais, there was still new cities reporting in the fourteenth week of 2021. Ceará and Rio de Janeiro are both big touristic states and also have a high air traffic flux of brazilian and foreign people, besides the reports of new cities were shifted in two weeks in relation to São Paulo, the dataset show that very few new cities reported cases after the first week of 2021. From the 5570 brazilian cities, the filtered dataset contains 2647 considering the ones that concentrated notifications and 5200 from the declared residence city.

To demonstrate the analysis involving filters by state, we chose the São Paulo state that obtained the highest number of records with a death outcome (126,702). In the top five states of this criteria are Rio de Janeiro (40,682) and Minas Gerais (35,201) that are also in Southeastern region. The other two states, Paraná (32,078) and Rio Grande do Sul (28,148), are located in the South region. The time-oriented analysis in weeks for this state indicate that from the pregnant female cases affected mainly those in the third and last quarter (58%). The overall cases in male (56%) were slightly higher than the one for females. In 2020 and 2021, the cases of the dataset were significantly higher for people whose age ranged from 41 to 70 (56.8%), followed by those with age above 71 (25.4%). The third more frequent age group was from 21 to 40 (15.7%), that surpassed the second group in the sixteenth week of 2021. The cases in white people represented 56% of the total, followed by the brown race (19%). The risk factors that most contributed in the overall cases was the asthma chronic condition that also affects the respiratory system (23.4% considering all cases, and 46% referring to the records with some risk factor). The second and third most frequent risk factors were kidney disease and systemic arterial hypertension. Still in São Paulo state, the analysis from the perspective of top ranked cities showed that São José do Rio preto (11,679 cases), Campinas (10,415), Ribeirão Preto (6,986) and São Bernado do Campo (6,466) contain the highest numbers of cases from people with risk factors. There were intervals of 11 days between the first symptoms and the notification date, but the most common delay to report was 7 (9% of cases) and 8 (8.2%) occurring mainly in the São Bernado do Campo, São José do Rio Preto e Campinas. In relation to the number of days passed between the notification to the outcome the most common values ranged form 3 to 6, corresponding together to 27.6% of the cases. Finally, the cities that received more people from other ones were São José do Rio Preto (7,218 cases), São Bernado do Campo (6,378) and Campinas (4,421).

### 3.2 Exploration of the mobility dataset

As for the srag analysis, the first verification analysis involve the assessment of data completeness and coverage for the mobility dataset. We checked the number of cities in each state that are contained in this dataset, and the Rio de Janeiro state has the highest coverage (93.5%), followed by Espírito Santo (76.92%) and São Paulo (64.19%). The missing data verification in this dataset refers to the empty values of change percentage observed for the six place categories, the states of the North and Northeastern regions in Brazil are the ones that have the highest mean percentage of empty values in these columns (ranging from 60% to 65%). For these states, the stratification of missing data percentage by place category shows that Grocery and farmacy, Retail and recreation and transit stations reaches more than 75%. In all states the workplace category has the lowest number of empty values (less than 5%). The correlation analysis (Figure 3) among the category places and the cases normalized per 1000 individuals demonstrates the positive strong correlation (96.1%) between the cases and the residential percentage change, this is expected since the adoption of lockdown was reactive so when the number of cases was exponentially increasing the people stayed at home, at the moment the cases decreased the restrictions were relaxed in most cities. The obvious strong negative correlation (ranging from -95.95% to -94.31%) among all other categories and the residential one was also expected because of their opposite nature. We chose the Rio de Janeiro state since this state obtained has more cities in this dataset, but these correlations were observed for all states in the same intensity (strong), and the time period was given in weeks.

**Fig. 3.**
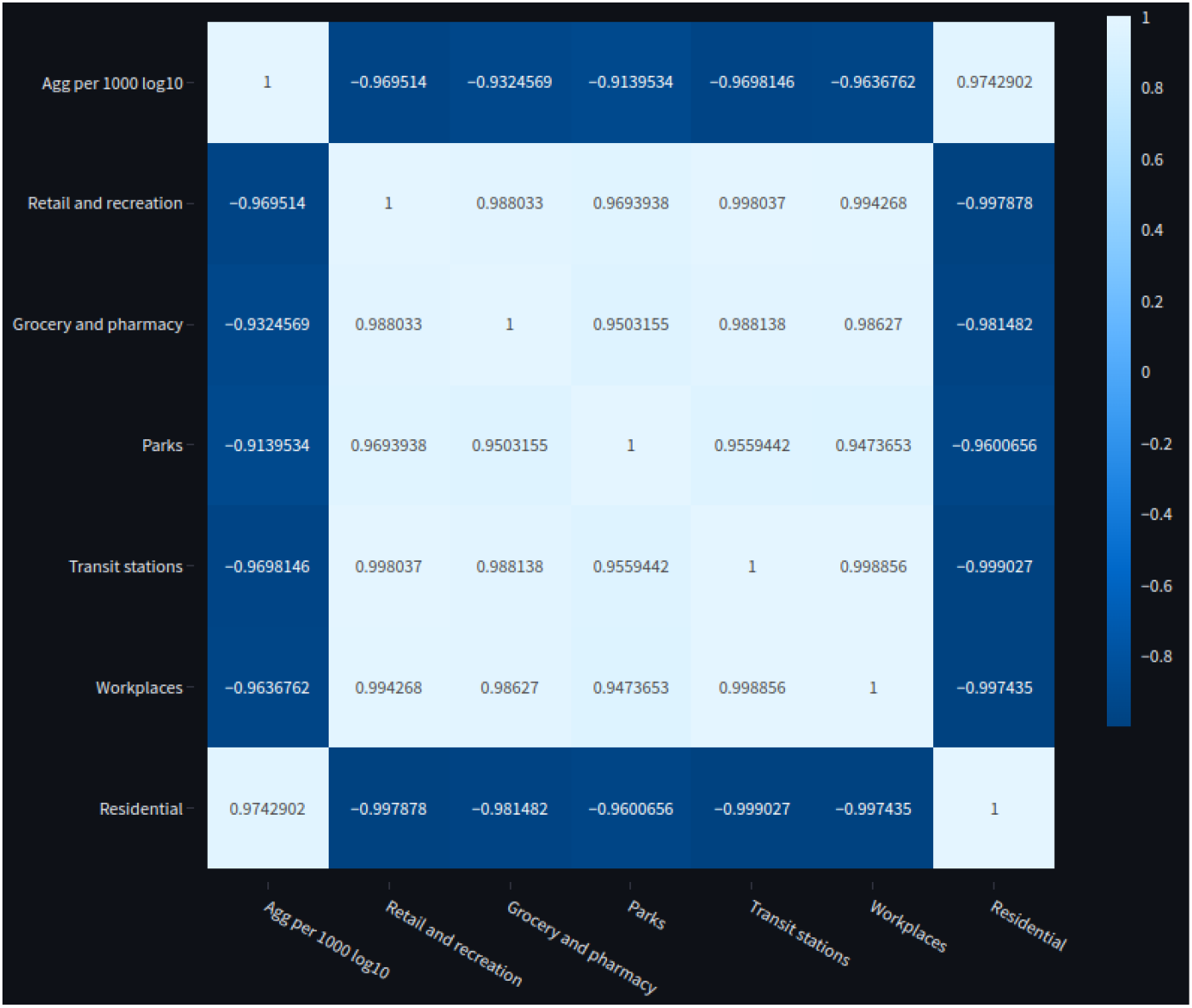
Correlation matrix among the six place categories and the number of cases normalized y population per 1000 in the Rio de Janeiro state.

The analysis of distribution (Figure 4) of residential and workplaces (the least empty column in the dataset) percentage changes along the cities for all states show that each state had a distinct behavior along their individual cities, there were outliers far from the overall mean for both categories in all states. In 12 states the overall median shows that there were no effect in the workplace mobility since they valued zero or close to zero. The Bahia state had the most negative outliers for workplaces and also positive ones for residential. The overall median fo residential mobility clearly shows that the lockdown was poorly implemented. This analysis only considered 2020 and 2021 years. Considering only the Rio de Janeiro state, the city that obtained the highest mean positive residential percentage change, along the observed period in weeks, was Itatiaia (15.2%), only other 12 cities in this state obtained a overall mean above 10%, which are: Santo Antônio de Pádua (14%), Armação dos Buzios (13.64%), Campos dos Goytacazes (13.30%), Niterói (12.57%), Mangaratiba (11.96%), Vassouras (11.25%), Macaé (11.07%), Paraíba do Sul (10.80%), Três Rios (10.41%), Rio das Ostras (10.31%), Rio de Janeiro (10.21%) and São João da Barra (10.02%).

**Fig. 4.**
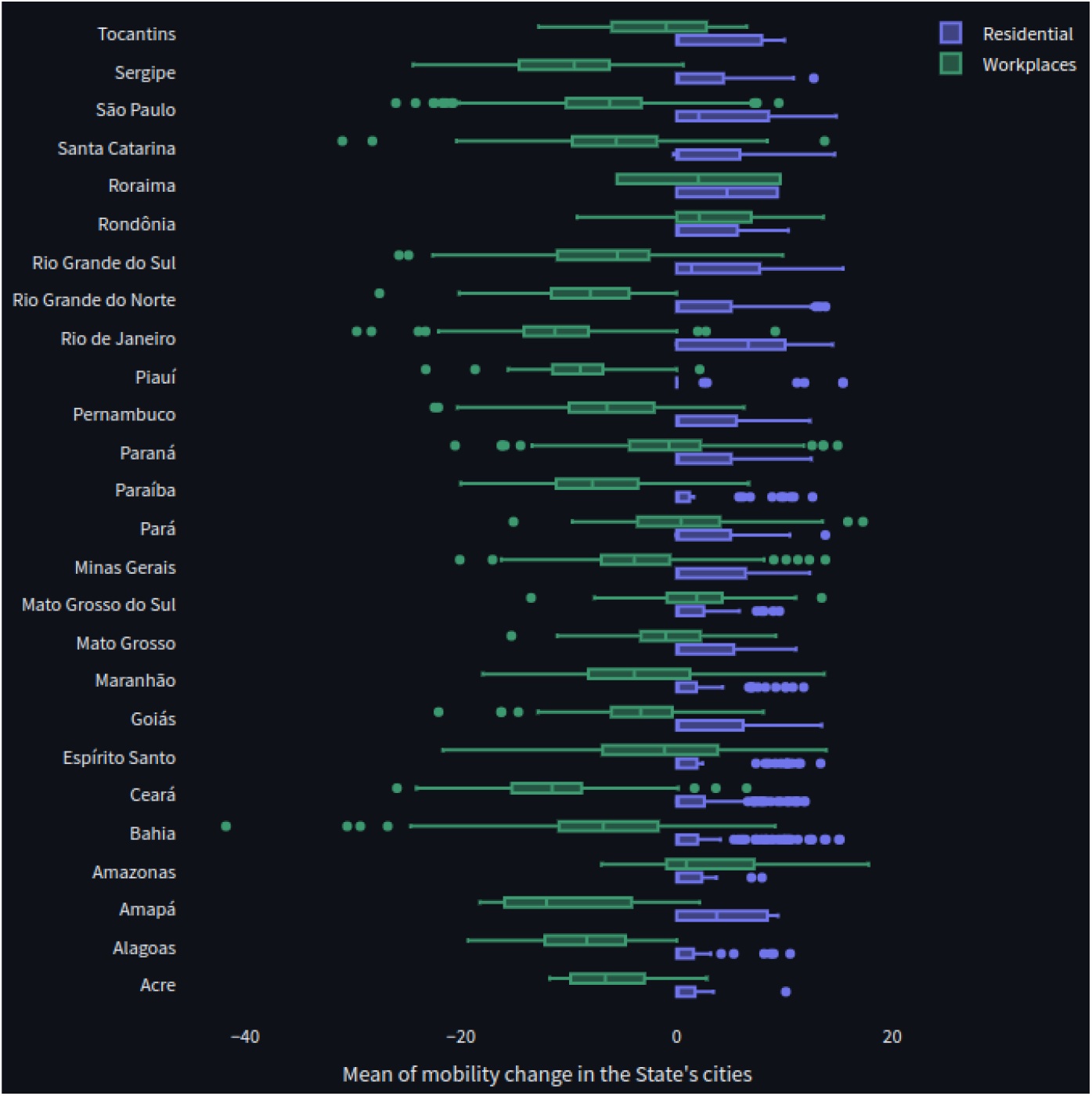
Dispersion of the overall means of percentage changes of residential and workplace categories in all cities available for each state.

### 3.3 City clustering

We performed an experiment to group cities according to their three features along the aggregated number of three features along weekly time intervals, which are the cases, deaths and the residential percentage change. This analysis had two goals: compare the clustering strategies by connected components detection [19] in a city network and by the traditional application of the K-means algorithm combined with the Elbow method [12]; Measure the agreement of the outcome-derived (cases and deaths) groups in relation to the residential feature from the mobility dataset. The input data corresponded to 127 numerical features for each of the 1810 cities that existed in srag and mobility datasets. The used the aggregated value per week scaled with base 10 log for the deaths and overall cases.

### 3.3.1 Clustering strategy comparison

Using the traditional k-means algorithm, the elbow technique found^10^ stopped in 13 groups for the overall cases, while the most optimized number of clusters for deaths and residential dimensions were 19 and 18, respectively. The distribution of cities along the clusters^11^ in the three versions was similar, with one cluster containing a minimum of 22 (residential), 23 (cases) and 32 (deaths), and the maximum number of elements in each group was 292 for the overall cases, 223 for residential and 199 for deaths. The mean number of elements was: 139 (Cases), 95 (Death) and 100 (residential). Besides the overall cases and deaths clusters diverge from each other in the number of clusters and cities arrangement, both share and agree with the classification of 23 capital cities in a same cluster (cl-9 for overall cases and cl-13 for deaths). In both, Palmas and Boa Vista were classified in another cluster (cl-10 for Cases and cl-3 for Deaths), the only difference was that while Macapá was in the same group as the majority of the capitals in the overall cases, this city was in a third distinct cluster for Deaths (cl-5). In relation to the residential percentage criteria, the major portion of the capitals (19) were grouped in cluster cl-12, Palmas was arranged with other common cities in cluster cl-0. The remaining capital cities (Macapá, Boa Vista, São Luís, Campo Grande, Maceió, Manaus) were arranged classified in the group cl-11.

The main difference between the two clustering strategies is that the traditional k-means will always classify all cities in some cluster, while the number of nodes (cities) in the network approach^12^ may vary according to the distance cutoff used to form the edges of the graph, low values of distance lead to smaller networks, since the temporal patterns of the three dimensions have distinct distributions. Using the distance of 10, the number of cities in the overall cases graph was 528, similar to the 523 for deaths, but both were smaller than the 653 in the residential graph. All the cities in the death graph were contained in the cases one, the five extra cities were Guará, Flórida Paulista, Guarulhos, Santo André and Barueri (all of them situated in São Paulo state). The cases graph shared 476 cities with the residential graph. The profile of the clusters using the network approach enlarge the number of detected clusters, decreasing their number of elements. The overall cases dimension obtained 149 groups, with a maximum of 27 cities in a cluster, a similar number was found for the death dimension with 147 groups and the same (27) maximum of cities. The residential dimension, generated more clusters (189) since there were more nodes in relation to the others, but the maximum number of cities was also 27. In all three graphs, most part of the groups contained 2 to 4 cities.

#### 3.3.2 Comparison among the overall cases and Death clusters against the residential derived groups

We computed the intersections among the clusters observed for cases and deaths with their closest corresponding groups in the residential set. This analysis checked the maximum correspondence found among the groups found using features of srag and mobility dataset. The higher the number of clusters, higher the possibility to find a cluster with the same members. This statement was confirmed comparing the number of cluster pairs among these datasets that obtained a sharing percentage of elements above 80%. Considering the groups generated by the k-means strategy^13^, only five Cases groups attended this criteria, whose intersection group elements varied from 18 (82% of agreement) to 90 (100%). Four clusters achieved more than 80% of consensus, whose number of elements in intersection ranged from 45 (100%) to 79 (96.34%). The number of cluster pairs increased significantly in the network strategy^14^, being 84 pairs for the Cases dimension and 85 for the death. Although most agreements achieved 100%, around 76.2% of the groups contained only 2 (54%) or 3 (22.6%) cities in both dimensions (cases and deaths).

### 3.4 Outcome prediction from mobility change features

We evaluated the capability of mobility percentage changes predicting the weekly aggregated values of each outcome (overall cases, deaths and cure). We fixed the formula the residential and tested it alone and combined in pairs with the other five place categories.

In general, for 98.47% of regression results^15^, the ordinary least square method used for regression returned the minimum values of root mean square errors for all cities, which indicates that the GLM models are more sensitive. We also measured the variation of rmse per method across cities grouped by brazilian regions, in the three outcomes the highest values (Table 3.4) were observed in the Center-West and South. Considering a cutoff of rmse under 0.005, the performance of the GLM predictor return 17 best ranked cities for overall cases, 32 for deaths and 34 for cures. The unique cities that were contained in the three ranked lists were: Vilhena located at the Rondonia state, with 95599 inhabitants; and Timon that belongs to he Maranhão state, with a population of 175044 inhabitants. From the oposite perspective the wors values of rmse observed were: 81.19 Cases) in Braço do Norte, 90.93 (Death) in Pirenópolis and 116.94 (Cure) in Santo Ângelo. Filtering the cities using a threshold of 0.5, the outcome prediction fits this restriction for 11% of the cities, while rising this threshold to 1 it reaches from 33.5% (Cases) to 39.7% (Death) of the cities. Using the portion of the dataset that attends the rmse value under 1, in the three outcomes there was a consensus that the co-variables that returned the major portion of these records were: residential + Grocery with pharmacy and residential + workplaces.

**Table 1.** Table comparing the rmse observed in the brazilian regions on predicting the overall cases, deaths and cures using the GLM and OLS regression methods.

| Methods<br>Region / Outcome | GLM |  |  | OLS |  |  |
| --- | --- | --- | --- | --- | --- | --- |
|  | Cases | Death | Cure | Cases | Death | Cure |
| **North** | 1.06 | 0.89 | 1.24 | 0.14 | 0.12 | 0.14 |
| **Northeast** | 0.80 | 0.70 | 0.72 | 0.16 | 0.13 | 0.15 |
| **Center-west** | 1.54 | 1.46 | 1.22 | 0.22 | 0.14 | 0.20 |
| **Southeast** | 0.80 | 0.66 | 0.73 | 0.20 | 0.15 | 0.18 |
| **South** | 1.67 | 1.51 | 1.58 | 0.20 | 0.15 | 0.18 |

The results considering the 26 capital cities, among the co-variables that led to the minor values of rmse for each outcome, the workplaces and grocery with pharmacy were the best combination with residential in the glm method while parks and workplaces ranked in the top for the ols. Ranking the capital results regardless of outcome and considering the only the minimum values of rmse, the top 50 records contain predominantly capitals from the northeastern and north regions, with occasional occurrences of São Paulo, Porto Alegre, Florianópolis, Cuiabá and Vitória. From the outcomes perspective, while the prediction of Cure and death obtained a rmse mean of 0.57, the cases obtained a higher rmse mean (0.63). We also observed that grocery with pharmacy and parks formed a consensus considering the top 2 co-variables in each outcome result groups.

### 3.5 Waves detection

This analysis allows the description and characterization of a series containing the overall case counts aggregated per time period and a window threshold. Our analysis only considered the detection of waves in the overall cases aggregated in weeks and the intervals among the neighbors in the waves should have at least a space of 3 (the window).

According to the results, 950 cities^16^ did not met the algorithm criteria to separate the waves, they should have a minimum number of cases and also follow a minimum homogeneous distribution of these cases along the weekly time series. From these 950 cities, on the one hand the maximum number of cases found was 1168 in a series whose reports were found only for 16 weeks, on the other hand some cities had 52 weeks filled with cases but the number of cases did not even reach 200.

The wave detection algorithm successfully found the cycles in 855 cities^17^, with the number of waves ranging from 1 to 8, but the major portion (85.73%) of cities obtained from 1 (30.87%) to 5 (11%) waves. Considering the capital cities, the total number of waves ranged from 3 (Belo Horizonte and Macapá) to 6 (Cuiabá, Florianópolis, Maceió and Salvador). According to the peaks computed from all waves, only five capitals reached a maximum value above 1000: Belo Horizonte (1476), Curitiba (1088), Recife (1452), Rio de Janeiro (1980) and São Paulo (4763).

## 4 Discussion

The dashboard web tool and the methods presented in this paper have as primary goal demonstrate the potential usage of each dataset for future pandemics. We focused in showing the covid-19 situation in Brazil according to the dataset information to describe the variety of forms to combine the features, derived columns and perspectives time and city oriented. The paper has no purpose to add epidemiological insights since the srag dataset coverage is far from the real number of cases and deaths.

The survInTime dashboard demonstrates visualization resources to explore open datasets by perspectives not elaborated by the published literature. [22] proposed a web tool to also explore the brazilian cases and perform some time series analysis. While they focused in epidemiological forecasting and other specific analysis, our dashboard proposes the exploration of new derived data and a more detailed and stratified analysis of the available columns in the srag dataset and its integration according to the time periods and cities with mobility changes in the six place categories. Our tool was not designed to perform epidemiological modeling.

All the methods demonstrated in this paper are publicly available in a Github repository^18^, with their respective documentation and usage instructions. The dashboard part of the repository contains all the processed data used for the analysis and the scripts to run the cases prediction from mobilility features, wave detection and city clustering are found in the separated folder *methods_ mlStats _analysis*^19^. In this folder, the user has also access to all the output data tables generated by the statistical and machine learning methods.

The two city clustering methods showed perspectives to analyze similar patterns among the cities in a using temporal aggregated features clustering. The strategies showed a consistent arrangement and agreement among each other, mainly concerning the capital cities. The network-based approach allows the user to prune and select the most significant nodes to build the graph using the distance threshold, while the traditional k-means approach provides an overview of the overall classification of the cities in an optimized number of groups. [23] also proposed a strategy to group brazilian cities but taking into consideration their relevant risk and other specific features of the cities such as social vulnerability, human development index, geographical area and other demographic factors. Our approach is designed to combine time-dependent evolution of features in two perspectives of filtering and analysis. Their strategy may be associated to enrich information about the cities that our approach grouped together. Our proposed strategy to predict overall cases, deaths and cures only from the google mobility changes observed from the place categories builds simple models but it attends the goal of attending a complete analysis stratification considering the brazilian cities, regions and the capitals behavior. We showed that besides the chaotic adoption and distribution of the lockdown intervals that affected the mobility in the cities, that were still the possibility of this kind of prediction in a portion of cities with a minimum error threshold. More sophisticated models have been proposed mixing models with other variables to enrich the formula given to the regression methods, such as vaccination and relative infectiousness and susceptibility [24]. [25] centered their experiment of outcomes prediction from the Google mobility dataset in the São Paulo city, studying the statistical distribution of the cases and deaths time series. They also prosed a residential mobility index (RMI) to track the isolation and exposure along the time. Finally, [26] tested the time series patterns of the google mobility dimensions, the covid-19 daily reports and the health departments using three strategies to detect seasonality in time series which are t-Distributed Stochastic Neighbor Embedding, Fast Fourier Transform, Seasonal Auto-Regressive Integrated Moving Average and transformers. This work only performed the analysis for the Araraquara city.

Finally, the last method proposed in this article describes the behavior of the overall cases time series behavior, returning the waves and their respective peak (maximum value, right before a drift) and the indexes of the series according to the trends of each step. The published literature concerning the study of waves intend to predict and forecast cases from them or detect their occurrences from other explainable variables [27–30]. Another perspective of wave analysis for covid-19 include the characterization of other aspects such as other diseases occurrences or clinical characteristics during the waves [31–35]. Our method intend to extract the isolated features of the cases time series and then enable multiple time series (multiple cities, for instance) characterization.

## Data availability

The data used in the manuscript are available online in a Github public repository at https://github.com/YasCoMa/dashboard-srag-mobility

## Code availability

The code is available in a Github public repository at https://github.com/YasCoMa/dashboard-srag-mobility

## Acknowledgments

Acknowledgments are not compulsory. Where included they should be brief. Grant or contribution numbers may be acknowledged.

## Footnotes

1 https://www.ibge.gov.br/estatisticas/sociais/populacao/22827-censo-demografico-2022.htm?edicao=35938&t=resultados

2 https://opendatasus.saude.gov.br/dataset/srag-2021-a-2023/resource/dd91a114-47a6-4f21-bcd5-86737d4fc734?inner_span=True

3 https://www.google.com/covid19/mobility/

4 https://opendatasus.saude.gov.br/dataset/srag-2021-a-2023/resource/dd91a114-47a6-4f21-bcd5-86737d4fc734?inner_span=True

5 https://www.google.com/covid19/mobility/

6 https://www.ibge.gov.br/estatisticas/sociais/populacao/22827-censo-demografico-2022.htm?edicao=35938&t=resultados

7 https://raw.githubusercontent.com/kelvins/Municipios-Brasileiros/main/json/estados.json

8 https://dashboard-srag-mobility-pipn9w3yh7p.streamlit.app/

9 https://www.reuters.com/graphics/world-coronavirus-tracker-and-maps/countries-and-territories/brazil/

10 https://github.com/YasCoMa/dashboard-srag-mobility/blob/master/methods_mlStats_analysis/results_clustering_elbow.tsv

11 https://github.com/YasCoMa/dashboard-srag-mobility/blob/master/methods_mlStats_analysis/results_clustering_kmeans.tsv

12 https://github.com/YasCoMa/dashboard-srag-mobility/blob/master/methods_mlStats_analysis/results_clustering_network.tsv

13 https://github.com/YasCoMa/dashboard-srag-mobility/blob/master/methods_mlStats_analysis/results_analysis_clustering_kmeans.tsv

14 https://github.com/YasCoMa/dashboard-srag-mobility/blob/master/methods_mlStats_analysis/results_analysis_clustering_network.tsv

15 https://github.com/YasCoMa/dashboard-srag-mobility/blob/master/methods_mlStats_analysis/results_regression_outcome_from_mobility.

16 https://github.com/YasCoMa/dashboard-srag-mobility/blob/master/methods_mlStats_analysis/not_detected_wave.tsv

17 https://github.com/YasCoMa/dashboard-srag-mobility/blob/master/methods_mlStats_analysis/waves_cities.tsv

18 https://github.com/YasCoMa/dashboard-srag-mobility/

19 https://github.com/YasCoMa/dashboard-srag-mobility/tree/master/methods_mlStats_analysis

## References

[1] Melo, C.M.L.D.E., Silva, G.A.S., Melo, A.R.S., Freitas, A.C.D.E.: COVID-19 pandemic outbreak: the brazilian reality from the first case to the collapse of health services. An. Acad. Bras. Cienc. 92(4), 20200709 (2020)

[2] Barberia, L.G., Costa, S.F., Sabino, E.C.: Brazil needs a coordinated and cooperative approach to tackle COVID-19. Nat. Med. 27(7), 1133–1134 (2021)

[3] Sansone, N.M.S., Boschiero, M.N., Marson, F.A.L.: Epidemiologic profile of severe acute respiratory infection in brazil during the COVID-19 pandemic: An epidemiological study. Front. Microbiol. 13, 911036 (2022)

[4] Lima, T.M., Palamim, C.V.C., Melani, V.F., Mendes, M.F., Pereira, L.R., Marson, F.A.L.: COVID-19 underreporting in brazil among patients with severe acute respiratory syndrome during the pandemic: An ecological study. Diagnostics (Basel) 12(6) (2022)

[5] Nascimento, I.J.B., Pinto, L.R., Fernandes, V.A., et al.: Clinical characteristics and outcomes among brazilian patients with severe acute respiratory syndrome coronavirus 2 infection: an observational retrospective …. Sao Paulo Med. J. (2020)

[6] Pereira, A.R., Branco, M.D.R.F.C., Costa, S.d.S.B., Lopes, D.A.M., Pinheiro, V.V., Oliveira, D.C.d., Pasklan, A.N.P., Gomes, J.A., Santos, A.M.D., Gama, M.E.A.: COVID-19 severe acute respiratory syndrome in brazilian newborns in 2020-2021. Rev. Bras. Epidemiol. 26, 230012 (2023)

[7] Godoi, A.P.N., Bernardes, G.C.S., Almeida, N.A.d., Melo, S.N.d., Belo, V.S., Nogueira, L.S., Pinheiro, M.d.B.: Severe acute respiratory syndrome by COVID-19 in pregnant and postpartum women. Rev. Bras. Saude Mater. Infant. 21, 461–469 (2021)

[8] Morais, R.B., Shimabukuro, P.M.S., Gonçalves, T.M., Hiraki, K.R.N., Braz-Silva, P.H., Giannecchini, S., To, K.K.W., Barbosa, D.A., Taminato, M.: Factors associated with death due to severe acute respiratory syndrome caused by influenza: Brazilian population study. J. Infect. Public Health 15(12), 1388–1393 (2022)

[9] Franco, V.F., Rodrigues, A.S., Junior, E.R.R., Godói, L.G., Monroy, N.A.J., Costa, R.A., Francisco, R.P.V.: Demographic and epidemiological characteristics of pregnant and postpartum women who died from severe acute respiratory syndrome in brazil: A retrospective cohort study comparing COVID-19 and nonspecific etiologic causes. PLoS One 17(10), 0274797 (2022)

[10] Sansone, N.M.S., Boschiero, M.N., Ortega, M.M., et al.: Severe acute respiratory syndrome by SARS-CoV-2 infection or other etiologic agents among brazilian indigenous population: an observational study from …. The Lancet Regional (2022)

[11] Delvenne, J.-C., Yaliraki, S.N., Barahona, M.: Stability of graph communities across time scales. Proc. Natl. Acad. Sci. U. S. A. 107(29), 12755–12760 (2010)

[12] Syakur, M.A., Khotimah, B.K., Rochman, E.M.S., Satoto, B.D.: Integration kmeans clustering method and elbow method for identification of the best customer profile cluster. In: IOP Conference Series: Materials Science and Engineering, vol. 336, p. 012017. iopscience.iop.org, ??? (2018)

[13] Sethuraman, N., Jeremiah, S.S., Ryo, A.: Interpreting diagnostic tests for SARS-CoV-2. JAMA 323(22), 2249–2251 (2020)

[14] Barabási, A.-L., Bonabeau, E.: Scale-free networks. Sci. Am. 288(5), 60–69 (2003)

[15] Craven, B.D., Islam, S.M.N.: Ordinary least-squares regression. The SAGE dictionary of quantitative (2011)

[16] Blume, J.D., Su, L., Olveda, R.M., McGarvey, S.T.: Statistical evidence for GLM regression parameters: a robust likelihood approach. Stat. Med. 26(15), 2919–2936 (2007)

[17] Chicco, D., Warrens, M.J., Jurman, G.: The coefficient of determination r-squared is more informative than SMAPE, MAE, MAPE, MSE and RMSE in regression analysis evaluation. PeerJ Comput Sci 7, 623 (2021)

[18] Iftimie, S., López-Azcona, A.F., Vallverdú, I., Hernández-Flix, S., Febrer, G., Parra, S., Hernández-Aguilera, A., Riu, F., Joven, J., Andreychuk, N., Baiges-Gaya, G., Ballester, F., Benavent, M., Burdeos, J., Catala, A., Castañé, E., Castañé, H., Colom, J., Feliu, M., Gabaldó, X., Garrido, D., Garrido, P., Gil, J., Guelbenzu, P., Lozano, C., Marimon, F., Pardo, P., Pujol, I., Rabassa, A., Revuelta, L., Ríos, M., Rius-Gordillo, N., Rodríguez-Tomas, E., Rojewski, W., Roquer-Fanlo, E., Sabaté, N., Teixidó, A., Vasco, C., Camps, J., Castro, A.: First and second waves of coronavirus disease-19: A comparative study in hospitalized patients in reus, spain. PLoS One 16(3), 0248029 (2021)

[19] He, L., Ren, X., Gao, Q., Zhao, X., Yao, B., Chao, Y.: The connected-component labeling problem: A review of state-of-the-art algorithms. Pattern Recognit. 70, 25–43 (2017)

[20] Ç etin, P., Amrahov, Ş.E.: A new network-based community detection algorithm for disjoint communities. TURK. J. OF ELECTR. ENG. COMPUT. SCI. 30(6), 2190–2205 (2022)

[21] Likas, A., Vlassis, N. J. Verbeek, J.: The global k-means clustering algorithm. Pattern Recognit. 36(2), 451–461 (2003)

[22] Marinho, P.R.D., Cordeiro, G.M., Coelho, H.F.C., Cabral, P.C.: The COVID-19 pandemic in brazil: Some aspects and tools. Epidemiologia (Basel) 2(3), 243–255 (2021)

[23] Martines, M.R., Ferreira, R.V., Toppa, R.H., Assunção, L.M., Desjardins, M.R., Delmelle, E.M.: Detecting space-time clusters of COVID-19 in brazil: mortality, inequality, socioeconomic vulnerability, and the relative risk of the disease in brazilian municipalities. J. Geogr. Syst. 23(1), 7–36 (2021)

[24] McBryde, E.S., Meehan, M.T., Adegboye, O.A., Adekunle, A.I., Caldwell, J.M., Pak, A., Rojas, D.P., Williams, B.M., Trauer, J.M.: Role of modelling in COVID-19 policy development. Paediatr. Respir. Rev. 35, 57–60 (2020)

[25] Ibarra-Espinosa, S., Freitas, E., Ropkins, K., Dominici, F., Rehbein, A.: Negative-Binomial and quasi-poisson regressions between COVID-19, mobility and environment in são paulo, brazil. Environ. Res. 204(Pt D), 112369 (2022)

[26] Aragão, D.P., Junior, A.G.d.S., Mondini, A., Distante, C., Gonçalves, L.M.G.: COVID-19 patterns in araraquara, brazil: A multimodal analysis. Int. J. Environ. Res. Public Health 20(6) (2023)

[27] Nesteruk, I.: Waves of COVID-19 pandemic. Detection and SIR simulations (2020)

[28] Soldi, G., Forti, N., Gaglione, D., Braca, P., Millefiori, L.M., Marano, S., Willett, P.K., Pattipati, K.R.: Quickest detection and forecast of pandemic outbreaks: Analysis of COVID-19 waves. IEEE Commun. Mag. 59(9), 16–22 (2021)

[29] Sîrbu, A., Barbieri, G., Faita, F., Ferragina, P., Gargani, L., Ghiadoni, L., Priami, C.: Early outcome detection for COVID-19 patients. Sci. Rep. 11(1), 18464 (2021)

[30] Yousefinaghani, S., Dara, R., Mubareka, S., Sharif, S.: Prediction of COVID-19 waves using social media and google search: A case study of the US and canada. Front Public Health 9, 656635 (2021)

[31] Souza, F.S.H., Hojo-Souza, N.S., Silva, C.M., Guidoni, D.L.: Second wave of COVID-19 in brazil: younger at higher risk. Eur. J. Epidemiol. 36(4), 441–443 (2021)

[32] Hoogenboom, W.S., Pham, A., Anand, H., Fleysher, R., Buczek, A., Soby, S., Mirhaji, P., Yee, J., Duong, T.Q.: Clinical characteristics of the first and second COVID-19 waves in the bronx, new york: A retrospective cohort study. Lancet Reg Health Am 3, 100041 (2021)

[33] Mitrofanova, L.B., Makarov, I.A., Gorshkov, A.N., Runov, A.L., Vonsky, M.S., Pisareva, M.M., Komissarov, A.B., Makarova, T.A., Li, Q., Karonova, T.L., Konradi, A.O., Shlaykhto, E.V.: Comparative study of the myocardium of patients from four COVID-19 waves. Diagnostics (Basel) 13(9) (2023)

[34] Silva, S.J.R.d., Pena, L.: Collapse of the public health system and the emergence of new variants during the second wave of the COVID-19 pandemic in brazil. One Health 13, 100287 (2021)

[35] Fu, R., Sutradhar, R., Li, Q., Hanna, T.P., Chan, K.K.W., Irish, J.C., Coburn, N., Hallet, J., Dare, A., Singh, S., Parmar, A., Earle, C.C., Lapointe-Shaw, L., Krzyzanowska, M.K., Finelli, A., Louie, A.V., Look Hong, N.J., Witterick, I.J., Mahar, A., Gomez, D., McIsaac, D.I., Enepekides, D., Urbach, D.R., Eskander, A.: Incident cancer detection during multiple waves of COVID-19: The tsunami after the earthquake. J. Natl. Compr. Canc. Netw. 20(11), 1190–1192 (2022)

